# Sleep Preserves, Wake Differentiates: Strength-Dependent Forgetting of Declarative Memories Across the Retention Interval

**DOI:** 10.1101/2025.11.03.686279

**Authors:** M. Schabus, M.S. Ameen, D.P.J. Heib

## Abstract

Sleep benefits memory, but whether this benefit depends on initial memory strength remains heavily debated. We asked whether sleep preferentially stabilizes weakly versus strongly encoded declarative memories, and whether classical sleep oscillations contribute to this effect.

Ninety healthy adults (n=30/group) learned 80 unrelated word pairs; half presented twice (weak, S-), and half three times (strong, S+). Participants were tested either immediately (∼40 min post-learning), after 9 hours containing nocturnal sleep monitored with polysomnography, or after a comparable daytime waking period. Memory performance was assessed using cued recall accuracy and reaction times (RT).

Strongly encoded items (S+) were recalled better than weakly encoded items (S−) across all groups. Overall memory performance was lowest after wakefulness, whereas performance after sleep was slightly higher than immediate recall. Crucially, wakefulness selectively impaired the consolidation of weak items, with the WAKE group forgetting disproportionately more S− than S+ items. In contrast, the SLEEP group retained weak and strong items similarly, yielding performance comparable to immediate recall. RT analyses showed generally slower responses for weak items, particularly after wakefulness, while response times in the sleep and immediate recall groups did not differ. Analyses of classical sleep features, such as slow oscillations density, sleep spindle intensity, and their coupling, revealed no significant associations with overnight retention. Correlations between these oscillations and memory performance were present both during the adaptation and the learning nights, suggesting trait-like, rather than state-dependent relationships. Overall, our findings indicate that sleep protects declarative memories irrespective of initial encoding strength.

## 1. Introduction

Sleep is widely recognized as beneficial for memory consolidation (Rasch & Born, 2013). Behaviorally, the most consistent finding is that sleep reduces time-dependent forgetting compared to wakefulness, an effect first noted by Ebbinghaus (1885) and is often attributed to reduced interference during sleep (Drosopolus et al., 2007; Ellenbogen et al., 2006; Müller & Pilzecker, 1900). Beyond this protective role, physiological processes have been proposed to actively support consolidation. Synaptic downscaling during deep sleep restores cortical homeostasis and may incidentally aid memory consolidation (Tononi & Cirelli, 2014), whereas the active systems consolidation framework highlights hippocampal reactivation and redistribution into cortical stores (Diekelmann & Born, 2010). Both views link consolidation to characteristic oscillations (slow oscillations and sleep spindles) which are thought to play a central role and may even act in complementary ways (Feld & Born, 2017).

Yet, the beneficial effects of sleep on memory do not seem to be uniform. Rather, several studies have reported sleep benefits only for subsets of the learned material, or particularly strong effects for specific types of memories. Sleep therefore appears to be selective, prioritizing the consolidation of certain memory traces over others depending on features such as salience or expected future relevance (Stickgold & Walker, 2013). Other than salience and future relevance, a range of candidate features have been discussed in this context, including emotional valence and, most fundamentally, initial memory strength. However, the literature remains inconsistent regarding the role of memory strength in sleep-dependent consolidation. For instance, some studies report that sleep preferentially stabilizes weakly encoded items (Drosopoulos et al., 2007; Denis et al., 2021), others conclude that strongly encoded items are favored for sleep-dependent consolidation (Tucker et al., 2008; Ellenbogen et al., 2009). Furthermore, Petzka et al. (2021) showed that under “standard” testing conditions without interference, a sleep-dependent consolidation effect is indeed only seen for weaker memories. Critically though, the authors report that with increased retrieval demands (i.e., learning new temporospatial arrangements of the same objects directly before final retrieval), sleep-dependent consolidation effects were seen for both weaker and stronger memories. These interesting results suggest that all memories may be consolidated during sleep, but that memories of different strengths benefit from sleep depending on the exact conditions before retrieval such as testing with or without retroactive interference before.

To complete the picture regarding the relevance of memory strength for sleep-dependent consolidation, Stickgold (2009, p. 306) proposed an inverted U-shaped curve of sleep benefit, suggesting that especially memories of intermediate strength profit from sleep, whereas extremely weak or strong items may not.

One possible reason for such divergent findings - besides the usual caveats of small sample sizes, varying tasks, and heterogeneous materials, which have hampered replication in the sleep-and-memory field (for a critical review see Nemeth et al., 2024) - is methodological. In many studies, memory strength is manipulated by a mixture of repeated study (“re-study”) and retrieval practice before sleep. Yet these two routes to strengthening are not equivalent: retrieval practice induces distinct neural responses (van den Broek et al., 2016), enhances retention more than re-study (Roediger & Butler, 2011), and in some cases even eliminates subsequent sleep effects (Bäuml et al., 2014). Importantly, later work including our own demonstrated that this pattern can be modulated by corrective feedback: when retrieval is followed by feedback, sleep-related benefits may re-emerge, particularly over longer retention intervals (Abel et al., 2019). Thus, retrieval and feedback jointly alter the mnemonic state of memories prior to sleep. Retrieval may thus constitute a form of consolidation, thereby diminishing or masking the additional role of sleep. This complicates the isolation of sleep’s direct contribution to memory stabilization.

In the present study, we therefore examined sleep’s impact on memory consolidation when initial strength is manipulated exclusively by re-study, without retrieval practice. That is, we omitted any pre-sleep testing, ensuring that all items entered the retention interval under comparable, retrieval-free conditions; encoding strength was manipulated through two versus three re-study sessions, creating weakly and strongly encoded items. We tested three independent groups: a sleep group with overnight retention, a wake group with equivalent daytime retention, and a short-delay group. The latter group not only serves as a control for circadian influences but also provides a benchmark for recall performance shortly after encoding, against which forgetting across the longer retention intervals (either filled with wake or sleep) can then be compared. Memory strength was assessed as (1) overall accessibility of memory traces (cued recall) and (2) the response time required for successful retrieval.

## 2. Methods

### 2.1. Participants

We recorded 92 participants. Two participants were excluded as their initial recall performance was too low (lower than 10%). The final sample included 90 healthy participants (70 females) aged between 18-30 years (M = 22.88, SD = 2.64). All participants were native German speakers, with no history of psychological or sleep disorders. We added the psychometric information about our participants in Suppl. Table 1. We divided the participants randomly into 3 experimental groups. The study was conducted in accordance with the ethical principles in the Declaration of Helsinki and was approved by the local research ethics committee (University of Salzburg Ethics Committee). All participants gave their written informed consent before the start of the experiment.

### 2.2. Experimental Design

The study employed a 3 x 2 mixed-factorial design, featuring a between-subjects factor GROUP with three levels (n=30/group): SHORT, SLEEP, and WAKE. The within-subjects factor of memory STRENGTH had two levels: weak (S-) and strong (S+).

Participants across all groups learned a list of word pairs before undergoing a retention interval, which involved either nocturnal sleep or daytime wakefulness, followed by a memory performance test (TEST). The SLEEP group studied the word pairs in the evening and completed the memory test the following morning after a 9-hour retention interval that included approximately 8 hours of sleep in the laboratory. Two or three days prior, participants completed an adaptation and screening night without performing the learning task. The WAKE group learned the word pairs in the morning and completed the memory test later the same evening after 9 hours of wakefulness. The SHORT group was tested 40 minutes post-encoding, matching the interval between the end of training and “lights off” in the sleep group, and thus served as a proxy for memory strength immediately prior to the sleep or wake retention interval (Figure 1). To control for potential circadian confounds, half of the subjects in the SHORT group encoded the word pairs in the evening, whereas the other half encoded them in the morning. Importantly, memory performance did not differ between evening-encoded (M = 74.3%) and morning-encoded items (M = 68.3%) within the SHORT group, F(1, 28) = .66, p = .20. Therefore, data from both subgroups were collapsed and treated as a single SHORT group in all subsequent analyses.

**Figure 1.**
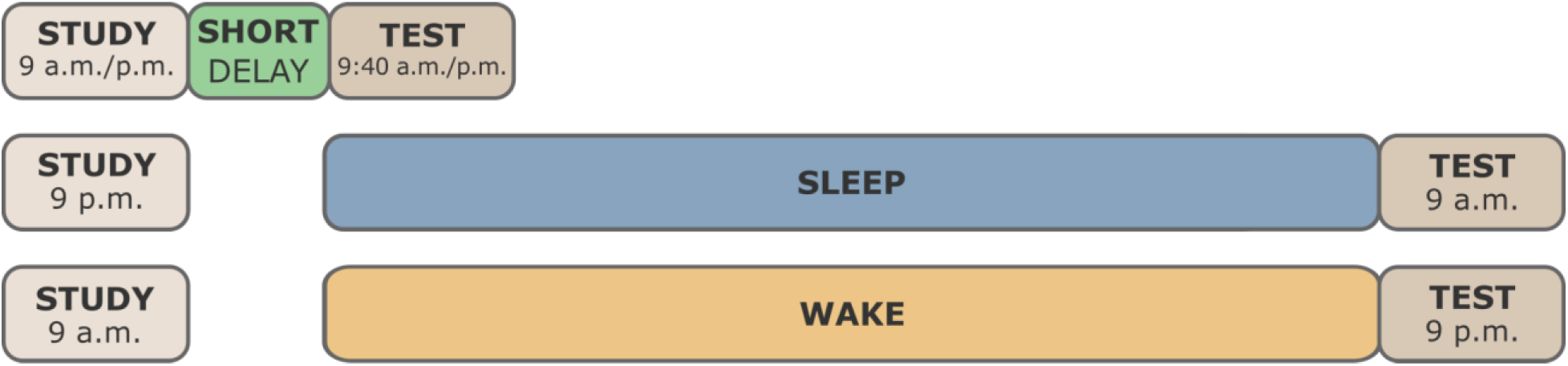
Experimental design. Following initial screening tests, 90 healthy participants were recruited to the experiment and were divided into 3 groups, SHORT, SLEEP and WAKE (N=30/group). Participants learned a list of unrelated word pairs and were tested after varying retention intervals: 9 hours containing nocturnal sleep (∼8h), 9 hours of wakefulness, or 40 minutes post-encoding. In total, 80 word-pairs were presented during the encoding phase, half of which (N=40) were presented twice (S-, weakly encoded) and the other half (N=40) was presented three times (S+, strongly encoded). Memory performance was assessed using a cued-recall test, measuring recall accuracy and response times for each word pair.

### 2.3. Word Pair Task

The study material consisted of a list of 80 unrelated word-pair associations. During the learning phase, half of the word pairs were presented twice (S-) and the other half three times (S+) (see Figure 2). Each word-pair was shown for 7500 ms, followed by a white fixation cross for 1500 ms and a red fixation cross for 1000 ms, indicating the end of encoding (white cross) and the start of the next word-pair presentation (red cross). The word-pair list was randomized for each participant, ensuring that identical word-pair repetitions were separated by the presentation of at least three other word pairs.

**Figure 2.**
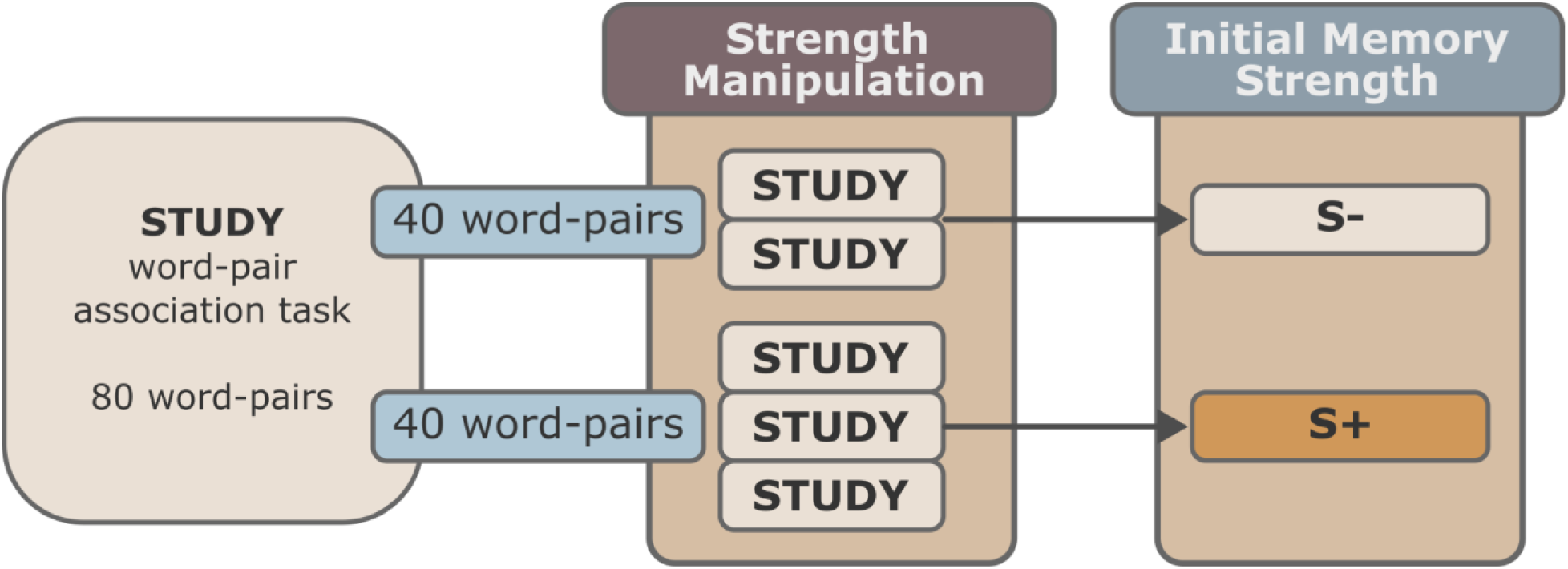
Word-pair learning task. Half of the word pairs were studied twice (S-) and the other half was studied three times (S+). Each word-pair was shown for 7500 ms, followed by a white fixation cross for 1500 ms, then a red fixation cross for 1000 ms, indicating the end of encoding (white cross) and the start of the next word-pair presentation (red cross). The word-pair list was randomized for each participant.

Memory performance was assessed using a cued-recall test, where participants were given the first word of each pair as a cue (for a maximum of 5000 ms) and asked to recall the corresponding second word. Participants were instructed to press a button on the keyboard as soon as they recalled the word, followed by their verbal response. Test performance was measured by recall rates (%) and reaction times.

### 2.4. EEG acquisition and analysis

EEG was recorded using Synamps EEG amplifiers (Neuro-Scan, Inc., El Paso, TX). EEG data were acquired using 16 Ag/AgCl electrodes (F7, F3, Fz, F4, F8, C5, C3, Cz, C4, C6, P3, Pz, P4, O1, Oz, and O2 as well as A1 and A2 for later re-referencing) attached to participants’ scalps according to the 10-20 system (Jasper, 1958). In addition, a number of physiological channels were recorded: one bipolar submental EMG channel, one bipolar electrocardiogram channel, one bipolar respiratory channel (chest wall movements), and two horizontal EOG channels. All channels were recorded using a sampling rate of 1000Hz, a high-pass filter (0.10 Hz) and a 50Hz notch filter. Monopolar channels were recorded using FCz as the online reference.

### 2.5. Sleep staging

Sleep was scored semi-automatically using an algorithm developed by the SIESTA Group (SIESTA GmbH) and described in (Anderer et al., 2005; 2010), following the standard criteria recommended by the American Association for Sleep Medicine (AASM; Berry et al., 2017).

### 2.6. Sleep spindle detection

Sleep spindles were detected using an algorithm developed by the SIESTA Group (AskAnalyzer; Gruber et al., 2015). First, we filtered the raw data between 11 and 16 Hz and then detected spindle events at electrodes re-referenced to the average of mastoids using the criteria described in Schimicek et al. (1994). Only events with an amplitude >12 µV and a duration between 500 and 2000 ms were considered. Further validation of the detected spindles was done using LDA in which the detected spindles were compared with a template that is generated based on the visual scoring of sleep spindles in 8730 min of PSG data from 189 healthy participants and 90 participants with sleep disorders. For our analyses, we detected sleep spindles at electrode C3 and only considered events that occurred during N2 and N3 sleep. We defined fast spindles as sleep spindles that occurred in the frequency range between 13-15 Hz, and we calculated sleep spindle intensity as described in Schabus et al. (2004), which is a measure quantifying the average of spindle duration x spindle amplitude.

### 2.7. Slow oscillation detection

To detect slow oscillations, we applied standard detection criteria (Mölle et al., 2004) using custom-built Matlab routines (MathWorks®, Natick, MA). The criteria were as follows: (1) a positive zero crossing follows a negative zero crossing within a time window of 0.3-1s; (2) the peak negativity between these zero crossings must exceed -80µV; and (3) the amplitude difference between the peak negativity and the subsequent positive peak must be at least 140µV. We measured slow oscillation density (at FCz) as the number of slow oscillations per minute in NREM (N2+N3).

### 2.8. Slow oscillation-spindle coupling

Slow oscillation-spindle coupling was assessed using an established individualized approach (Helfrich et al., 2018; Hahn et al., 2020). Each participant’s sleep spindle peak frequency during NREM N2 and N3 sleep was identified from the oscillatory component of EEG power spectra computed from 15-s epochs, providing a frequency resolution of 0.067 Hz. The spindle peak was defined as the largest oscillatory peak between 10 and 17 Hz, averaged across frontal, central, and occipital electrodes. Spindles and slow oscillations were detected automatically for each channel.

Spindles were identified by bandpass filtering the signal within ±2 Hz of the individual spindle peak, followed by Hilbert transformation and amplitude thresholding (75th percentile, duration 0.5-3 s). Slow oscillations were detected from data filtered from 0.3 to 2 Hz using zero crossings, with cycle durations of 0.8-2 s and amplitudes exceeding the 75th percentile. Events were epoched from -2.5 to 2.5 s around slow oscillations’ troughs or spindle peaks, and artefactual epochs were excluded.

Slow oscillation-spindle coupling was quantified using an event-locked cross-frequency coupling approach. After Z-transformation, slow oscillations’ phase was extracted at the time of maximal spindle amplitude. Analyses were restricted to slow oscillations epochs containing a spindle within ±1.5 s of the slow oscillation trough. Coupling strength was quantified using circular statistics as mean vector length (MVL). Event-locked time-frequency representations centered on slow oscillation troughs were computed using a 500-ms Hanning window (5-30 Hz, 0.5 Hz resolution), baseline-corrected, and z-transformed using a bootstrapped baseline (5000 iterations, -2 to -1.5 s). Due to missing or corrupted sleep data from at least one recording night, seven participants were excluded, resulting in a final sample of 23 participants for this analysis.

### 2.9. Statistical analyses

Statistical analyses were conducted in R, using a 3 (Group: SHORT, SLEEP, WAKE) × 2 (Condition: S−, S+) mixed-design ANOVA, with Group as a between-subjects factor and Condition as a within-subjects factor. The dependent variables were memory performance, operationalized as the percentage of correctly recalled items, and response time, defined as the latency between cue-word onset and the participant’s button press indicating successful retrieval. Effect sizes are reported as generalized eta squared (η^2^_G). Where appropriate, post hoc comparisons were performed using Bonferroni-corrected pairwise t-tests. Assumptions of normality, homogeneity of variance, and homogeneity of covariance were assessed and were deemed acceptable given the balanced design and the robustness of ANOVA to moderate violations. For statistics on reaction times, we removed subjects with less than 10 available reaction times in either condition from the analyses as we aimed for reliable reaction time estimates per subject.

Associations between physiological sleep measures and memory performance were assessed using Spearman rank correlations throughout. Spearman correlations were chosen as a conservative, distribution-independent measure that is well suited for relating behavioral performance to physiological sleep parameters. To compare correlation coefficients between conditions (S− vs. S+), we used a percentile bootstrap method for paired samples, implemented via the twoDcorR function from Wilcox’s robust statistics toolbox (Rallfun-v35), following the procedure described by Wilcox and Muska (2002). This approach allows for direct statistical comparison of dependent correlation coefficients without relying on strict parametric assumptions. Thus, correlation analyses were hypothesis-driven and limited to a small set of predefined sleep metrics; no correction for multiple comparisons was applied, and no statistically significant effects are claimed.

For slow oscillation-spindle coupling, to compare coupling strength between Adaptation and Experimental nights, data were first assessed for normality using the Shapiro-Wilk test applied independently to each condition. If both conditions met the normality assumption (p > 0.05), paired-samples t-tests were used; otherwise, Wilcoxon signed-rank tests were applied. For each electrode, mean coupling strength values were computed for both conditions, and test statistics and p-values were extracted. To control for multiple comparisons across electrodes, p-values were adjusted using the false discovery rate (FDR) procedure. For correlations between coupling strength during experimental night and percentage of recalled words, normality of both neural and behavioral variables was assessed using Shapiro-Wilk tests. Consequently, Spearman’s correlations were used for all analyses. Correlations were calculated separately for strongly encoded (S+) and weakly encoded (S-) items and for each electrode. Resulting p-values were corrected for multiple comparisons using FDR adjustment.

## 3. Results

In this section, we present the behavioral memory performance data as well as the association with slow oscillations, sleep spindles and (for the sleeping group). Demographic sleep data are reported in Suppl. Table 2.

### 3.1. Memory performance

The analysis revealed a strong main effect of STRENGTH, F(1, 87) = 172.87, p < .001, η^2^_G = .17, indicating higher recall performance for S+ than for S− items across all groups. In addition, there was a robust main effect of GROUP, F(2, 87) = 28.99, p < .001, η^2^_G = .37, reflecting overall differences in memory performance between the SHORT, SLEEP, and WAKE groups (Figure 3, left). Importantly, these main effects were qualified by a significant but comparatively small GROUP × STRENGTH interaction, F(2, 87) = 5.75, p = .005, η^2^_G = .014, indicating that the magnitude of the strength effect differed across retention conditions.

**Figure 3.**
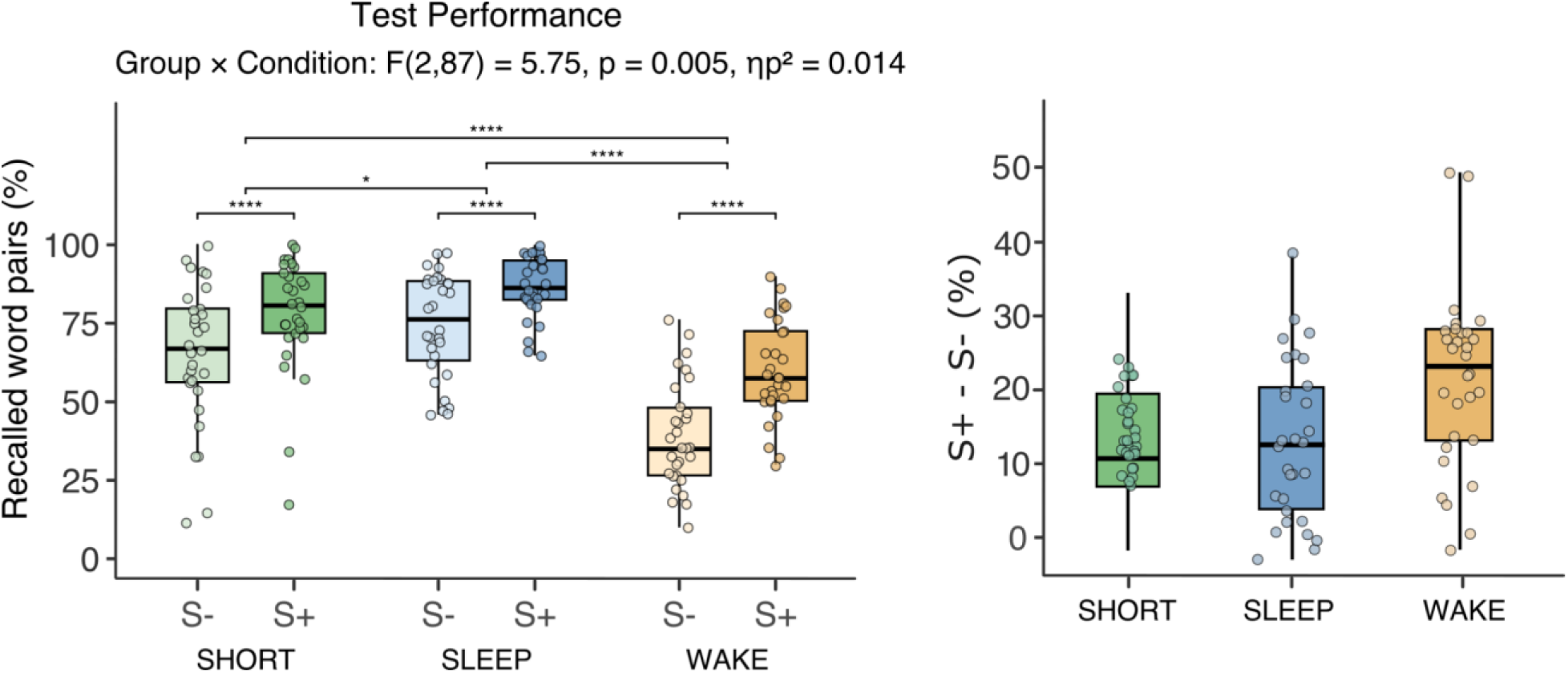
Behavioral performance for correctly retrieved items (HITS). Displayed are the 3 groups with immediate recall (SHORT), overnight sleep (SLEEP) and WAKE between learning and final retrieval. S-denotes items that were studied twice whereas S+ denotes items studied thrice in the encoding face. Note that the SLEEP and WAKE groups did not have a recall following immediately upon learning and reveals that S+ items are always superior in recall to S-items independent of the group assignment. Sleep group performance is identical to immediate recall after learning whereas the wake group demonstrates behavioral decrements with respect to both immediate recall and sleeping after learning. Sleep-dependent consolidation mechanisms.

Post-hoc pairwise comparisons confirmed that S+ items were recalled significantly better than S− items in all three groups (all ps < .001). Mean performance levels were as follows: SHORTS+ = 77.8% (SD = 18.2), SHORTS-= 65.2% (SD = 22.5); SLEEPS+ = 86.6% (SD = 9.8), SLEEPS-= 74.3% (SD = 16.6); WAKES+ = 60.0% (SD = 15.8), WAKES-= 39.2% (SD = 16.9).

Collapsing across strength levels, overall memory performance differed reliably between groups. Participants in the WAKE group performed significantly worse than both the SLEEP group (p < .001) and the SHORT group (p < .001). Importantly, the SLEEP group also outperformed the SHORT group (p = .028), indicating an overall benefit of sleep on memory performance relative to immediate testing.

Inspection of the GROUP × STRENGTH interaction suggested that this effect was driven by a disproportionate loss of weakly encoded (S−) items during wakefulness. While the strength effect (S+ > S−) was present in all groups, the performance gap between S+ and S− items was comparable in the SHORT (Δ ≈ 12.6 percentage points) and SLEEP groups (Δ ≈ 12.3 percentage points), but substantially larger in the WAKE group (Δ ≈ 20.8 percentage points) (Figure 3, right).

To further clarify the nature of this interaction, we conducted an additional analysis quantifying memory change relative to immediate performance. Specifically, performance in the SLEEP and WAKE groups was expressed as the difference from the corresponding SHORT group baseline (with evening-tested SHORT participants serving as the reference for the SLEEP group and morning-tested SHORT participants for the WAKE group), yielding an estimate of forgetting or gain across the 9-hour retention interval. This follow-up mixed ANOVA with GROUP (SLEEP vs. WAKE) and STRENGTH (S−, S+) revealed a large main effect of GROUP, F(1, 58) = 47.89, p < .001, η^2^_G = .41), indicating substantially greater forgetting following wakefulness than sleep. There was also a main effect of STRENGTH, F(1, 58) = 6.83, p = .011, η^2^_G = .017), as well as a significant GROUP × STRENGTH interaction, F(1, 58) = 7.17, p = .010, η^2^_G = .017).

Follow-up comparisons (Figure 4) showed no difference in forgetting between S− and S+ items in the SLEEP group, t(29) = .05, p = .962, indicating comparable retention of weak and strong memories across sleep (S−: Δ ≈ +6.2 percentage points; S+: Δ ≈ +6.1 percentage points relative to immediate recall). In contrast, the WAKE group exhibited significantly greater forgetting for S− than for S+ items, t(29) = −3.56, p = .001, with weakly encoded items showing a substantially larger performance decline (S−: Δ ≈ −22.6 percentage points) than strongly encoded items (S+: Δ ≈ −14.8 percentage points).

**Figure 4.**
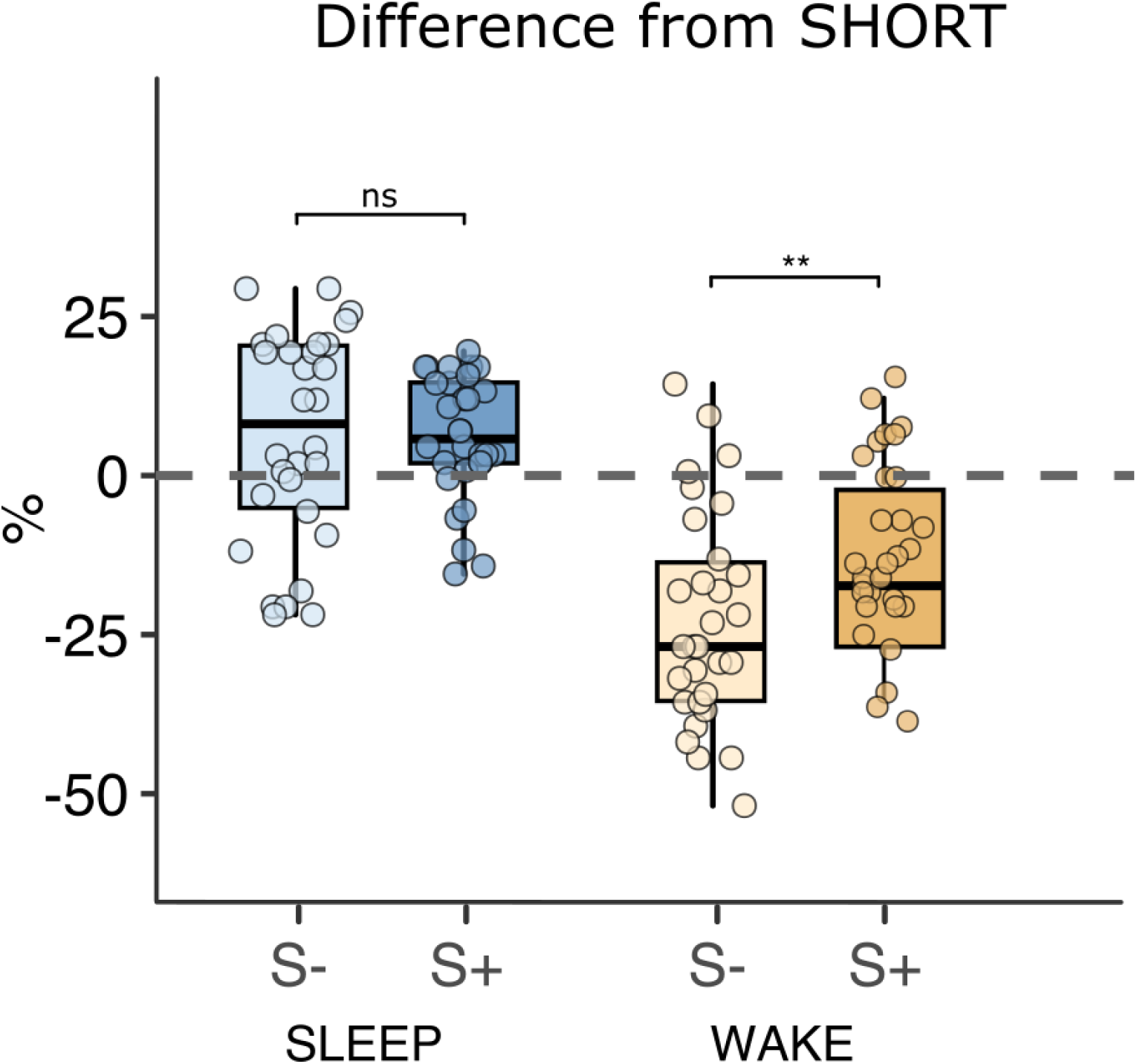
Change in memory recall relative to immediate performance in the SLEEP and WAKE groups. In the SLEEP group, post-retention recall did not differ between strongly (S+) and weakly (S−) encoded items. In contrast, in the WAKE group, weakly encoded items (S−) exhibited significantly greater forgetting than strongly encoded items (S+).

Reaction time data were analyzed using the same mixed 3 × 2 ANOVA with GROUP (SHORT, SLEEP, WAKE) as a between-subjects factor and STRENGTH (S−, S+) as a within-subjects factor. The analysis revealed a significant main effect of STRENGTH, F(1, 77) = 57.04, p < .001, η^2^_G = .07, indicating slower responses for weakly encoded (S−) compared with strongly encoded (S+) items. In addition, there was a significant main effect of GROUP, F(2, 77) = 12.86, p < .001, η^2^_G = .23. Post-hoc comparisons showed that participants in the WAKE group responded significantly more slowly than those in both the SHORT and SLEEP groups (both ps < .001), whereas response times did not differ between the SHORT and SLEEP groups (p = 1.00). Mean response times were 1611 ms (SD = 295) for the SHORT group, 1660 ms (SD = 359) for the SLEEP group, and 1998 ms (SD = 275) for the WAKE group (Figure 5).

**Figure 5.**
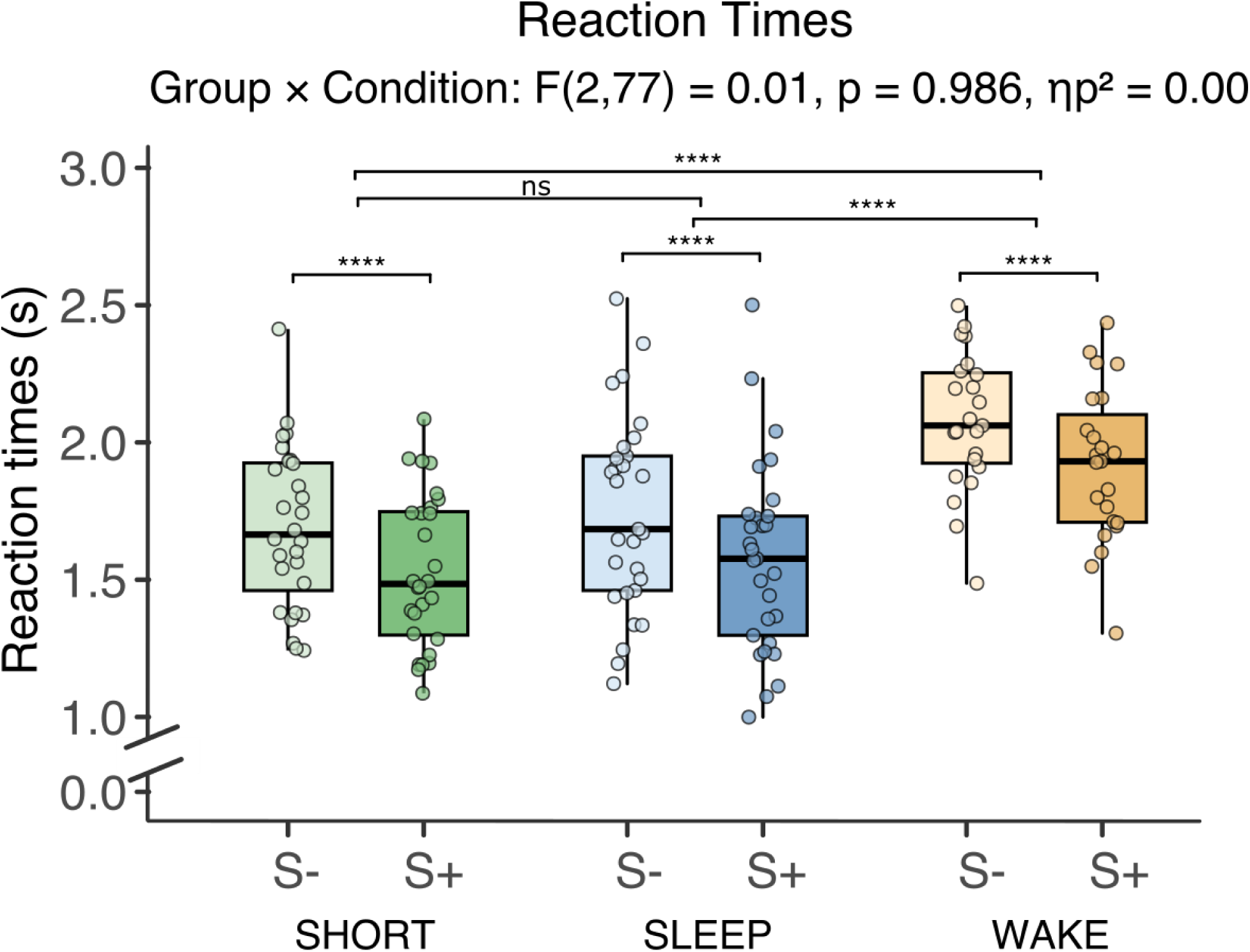
Reaction times for correctly retrieved items (HITS). Displayed are the 3 groups with immediate recall (SHORT), overnight sleep (SLEEP) and WAKE between learning and final retrieval. S-denotes items that were studied twice whereas S+ denotes items studied thrice in the encoding face. Note that the SLEEP and WAKE groups did not have a recall following immediately upon learning and reveals that S+ items are always superior in recall to S-items independent of the group assignment. Sleep group performance is identical to immediate recall after learning whereas the wake group demonstrates slower reaction times as compared to SHORT and SLEEP groups.

Importantly, the GROUP × STRENGTH interaction was not significant, F(2, 77) = 0.01, p = .90, η^2^_G < .001, indicating that the slowing observed after wakefulness did not differentially affect weakly versus strongly encoded items.

### 3.2. Influence of Sleep microstructure on memory performance

For the 30 participants in the sleep group, we additionally correlated the recall performance of S-and S+ items with classical metrics that have been associated with declarative learning in the past. We focused on slow oscillation density (at FCz) as well as fast sleep spindle intensity (13-15Hz spindle amplitude x duration at C3).

Spearman correlations (2-tailed) revealed that fast spindle intensity showed non-significant positive correlations with recall performance of S- and S+ items (for S-items: r_27_= .25, p= .20 in the adaptation night; r_28_= .22, p= .24 in the learning night). For S+ items, positive correlations were revealed in the adaptation (r_27_= .39, p= .04) as well as the learning night (r_28_= .37, p= .04) (cf. Figure 6, upper panels). Analysis reveals that correlations of the adaptation nights (p= .33) as well as the experimental learning nights (p= .30) do not statistically differ. These findings indicate that learning before sleep did not alter the association of studied sleep metrics and memory performance. Similarly, the numerical stronger correlations in the learning as compared to the adaptation night are not statistically different for S- (p= .82) as well as S+ (p= .82) items.

**Figure 6.**
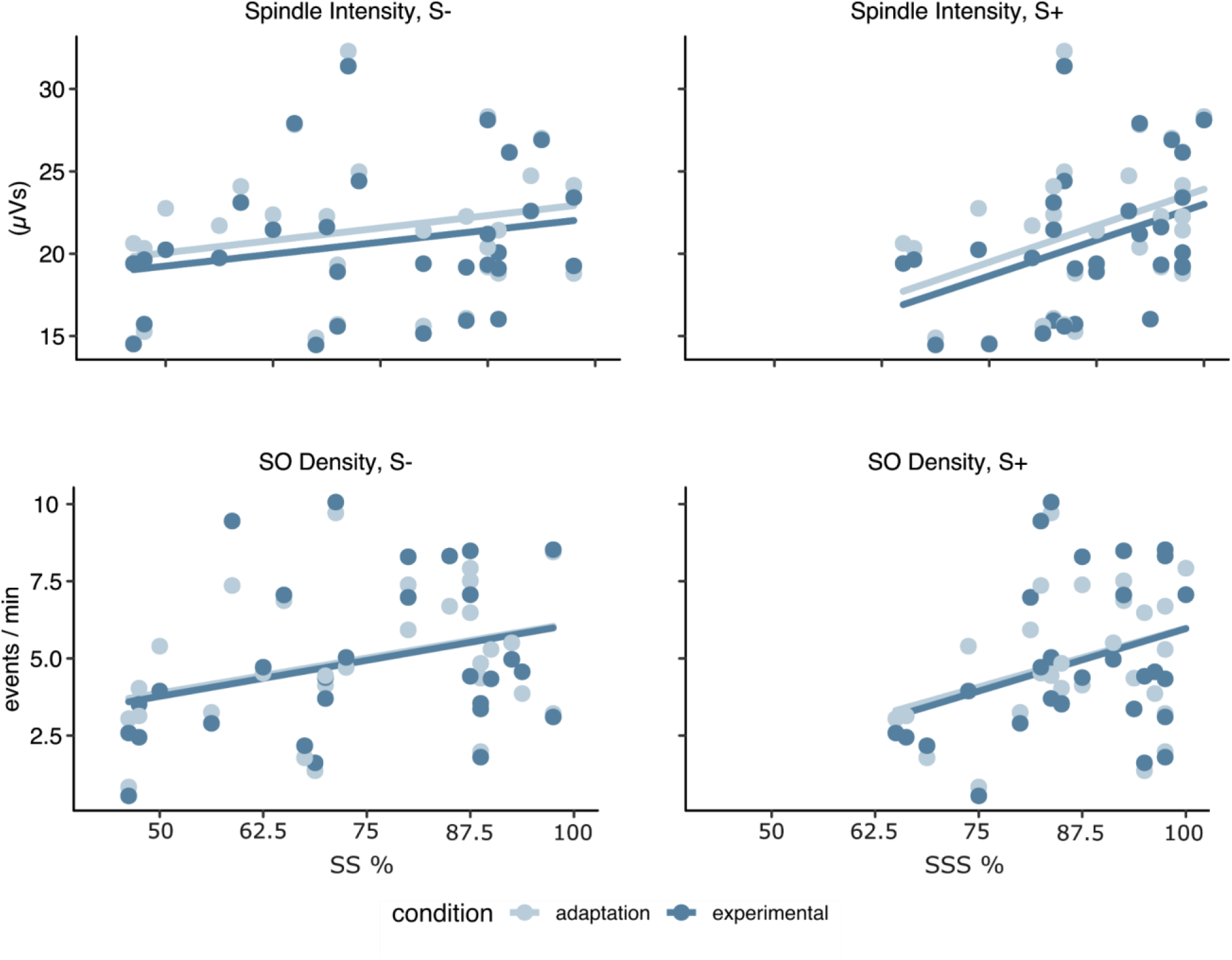
Correlations of fast spindle intensity and slow oscillation density with recall performance (sleep group). The upper left panel depicts the correlation between recall performance of weakly encoded items (S-) with spindle intensity in the adaptation (light blue color) as well as the learning night (dark blue color), with the right panel highlighting the associations for S+ items. The lower two panels illustrate positive associations of weak S- and strong S+ items with slow oscillation density. Note that these associations are already present in the non-learning adaptation nights and are statistically not different from the association in the learning night.

Spearman correlations (2-tailed) showed positive but non-significant associations between slow oscillation density and recall performance for both weak (S−) and strong (S+) items. For S− items, recall performance correlated positively with slow oscillation density in both the adaptation night (r(27) = .33, p = .08) and the learning night (r(27) = .30, p = .12). Similarly, for S+ items, positive but non-significant correlations were observed in the adaptation night (r(27) = .34, p = .11) and the learning night (r(27) = .27, p = .16) (see Figure 6, lower panel). Importantly, the strength of the correlations did not differ significantly between the adaptation and learning nights, neither for S− items (p = .77) nor for S+ items (p = .62).

Additionally, we explored the role of Slow oscillation-spindle coupling in memory performance. Our results indicated that i) there was no significant change in slow oscillation-spindle coupling between the adaptation and experimental nights (see Suppl. Figure 1 and Suppl. Table 3 for statistics), and ii) no significant correlation between slow oscillation-spindle coupling during the experimental night and overnight memory performance (see Suppl. Figures 2-3 and Suppl. Table 4 for statistics).

## 4. Discussion

Our findings show that sleep prevents the forgetting of declarative memories regardless of encoding strength, with slightly more pronounced forgetting of weak items during wakefulness. Memory retention differed between groups, with superior performance following sleep compared to wakefulness. When collapsing across encoding strength, retention after sleep was also slightly higher than immediate recall. This behavioral benefit was not modulated by memory strength, as weakly and strongly encoded items were retained to a similar extent after sleep, whereas wakefulness was associated with disproportionate forgetting of weakly encoded items. Analyses of canonical sleep microevents provided no converging mechanistic support: neither fast spindle density, slow oscillation density, nor their temporal coupling showed stronger learning-night effects (relative to the adaptation night), and none of these metrics showed robust associations with overnight retention performance.

A central methodological feature of the present study was the omission of an immediate recall test in the SLEEP and WAKE groups. Instead, a separate SHORT group provided an immediate estimate of post-encoding baseline performance, avoiding pre-retention retrieval, which is known to act as a potent mnemonic manipulation that can alter subsequent sleep-dependent consolidation (Bäuml et al., 2014; Abel et al., 2019). This design ensured that memory strength was manipulated exclusively via re-exposure (S− vs. S+) and that all items entered the retention interval under comparable, retrieval-free conditions.

By manipulating encoding strength, Petzka et al. (2021) showed that under standard testing conditions, sleep benefits primarily favor weak memories, whereas both weak (S−) and strong (S+) memories benefit when retrieval is challenged by interference. Our results align with this pattern: under standard retrieval without interference, sleep preserved both weak and strong memories at post-encoding levels, while wakefulness disproportionately impaired weak items. This pattern poses sleep as a primarily protective, rather than selectively enhancing, state in the present paradigm. On a similar note, it should be noted that our binary manipulation of encoding strength may not sufficiently sample the full continuum required to test the inverted U-shaped consolidation account proposed by Stickgold (2009). Thus, the absence of a strength-dependent sleep effect in the present study should not be taken as evidence against such non-linear relationships but rather suggests that within the strength range examined here, sleep did not differentially benefit weak versus strong memories.

In our paradigm, all sleep measures showed predominantly positive, albeit non-significant, correlations with overnight consolidation. Importantly, these associations were already present during the non-learning adaptation night and did not differ between nights. This suggests that the observed relationships are not specific to memory consolidation processes. These traditional sleep metrics therefore did not predict overnight retention performance but rather reflect a general trait-like relationship (Bódizs et al., 2005; Schabus et al., 2006) whereby individuals with better memory performance tend to have more sleep spindle activity (e.g., Wislowska et al., 2016) and/or slow oscillations (e.g., Qin et al., 2025).

We also found no evidence that slow oscillation-spindle coupling strength differed between the learning night and the non-learning adaptation night, given prior work emphasizing precise spindle nesting within slow oscillation upstates as a key mechanism of sleep-dependent memory consolidation (Helfrich et al., 2018; Klinzing et al., 2019; Staresina et al., 2015). Moreover, coupling strength in the learning experimental night did not correlate with overnight retention performance for either weakly or strongly encoded items. These findings suggest that, at least in this paradigm, slow oscillation-spindle coordination does not index learning-specific consolidation processes. Instead, the absence of learning-related modulation, together with similar associations across adaptation and learning nights, points toward more trait-like relationships between sleep microarchitecture and memory performance. While a substantial body of evidence supports a role for slow oscillation-spindle coupling in sleep-associated memory consolidation (see meta-analysis by Ng et al., 2025), our data suggest that this mechanism is not universal. Instead, its contribution may depend on task demands, initial encoding strength, and retrieval conditions.

From a theoretical perspective, our findings align more closely with permissive or protective models of sleep-dependent memory consolidation than with accounts emphasizing an active, experience-dependent role of sleep. Memory performance after nocturnal sleep was statistically indistinguishable from immediate morning recall, suggesting that sleep primarily stabilized memories at their post-encoding level rather than enhancing them beyond baseline. This pattern is consistent with the idea that sleep reduces retroactive interference by shielding memories from waking input, as proposed by permissive consolidation frameworks (Diekelmann & Born, 2007). In contrast, we found little evidence supporting an active consolidation account in which learning-specific sleep oscillations drive memory strengthening. Classical sleep microevents such as slow oscillations and fast spindles did not correlate more strongly with retention following the learning night compared to a non-learning adaptation night, and these correlations did not differ statistically between nights. Moreover, fast spindle density was not increased following learning demands but instead showed a numerical decrease from adaptation to experimental nights. Together, these findings argue against a learning-induced upregulation of oscillatory mechanisms supporting active replay, as suggested by systems consolidation models (Diekelmann & Born, 2010; Klinzing et al., 2019).

These findings indicate that sleep’s impact on retention, at least in the used declarative word-pair paradigm, is not modulated by nor selectively tuned to the strength of the memory engrams before falling asleep. Since memory strength exists on a continuous spectrum, future work may need to probe a wider range of encoding conditions, strength manipulations of items before sleep, as well as manipulations to challenge the memory traces before retrieval such as interference tasks before retrieval following sleep. Additionally, in this study, we only measured memory after 9 hours following initial encoding but not at later time points where effects could vary dramatically. Moreover, we only focused on the “usual suspects” that were earlier associated with sleep-dependent declarative memory consolidation; such as slow oscillations and sleep spindles. It is possible that other sleep markers are better at indexing the mechanisms needed to strengthen declarative memories at night.

Future research should aim to move beyond the classical oscillatory markers examined here and explore a broader spectrum of neural and physiological signatures that may underlie the protective effects of sleep on memory. High-density EEG, intracranial recordings, or multimodal imaging could reveal finer-grained interactions between hippocampal and cortical networks that remain hidden in conventional scalp measures. Moreover, expanding the temporal window beyond a single night, particularly through continuous, real-world monitoring approaches, as for example demonstrated by Topalidis et al. (2023), may uncover delayed or cumulative effects of sleep on differently encoded memories, offering insights into how consolidation unfolds dynamically across multiple nights. The demonstrated feasibility of accurate daily sleep staging using low-cost wearable devices opens the door for large-scale, longitudinal studies linking nightly physiological fluctuations to memory outcomes. Experimental manipulations of interference, reactivation, and targeted stimulation could further clarify whether sleep’s stabilizing role is merely passive protection or reflects active integration of memories into long-term memory networks. Ultimately, understanding why sleep preserves all memories equally in some contexts, but selectively in others, will help refine theoretical models of memory consolidation and bridge trait-like and state-dependent perspectives on how the sleeping brain shapes our cognitive architecture.

## 6. Appendix

**Supplementary Table 1.**
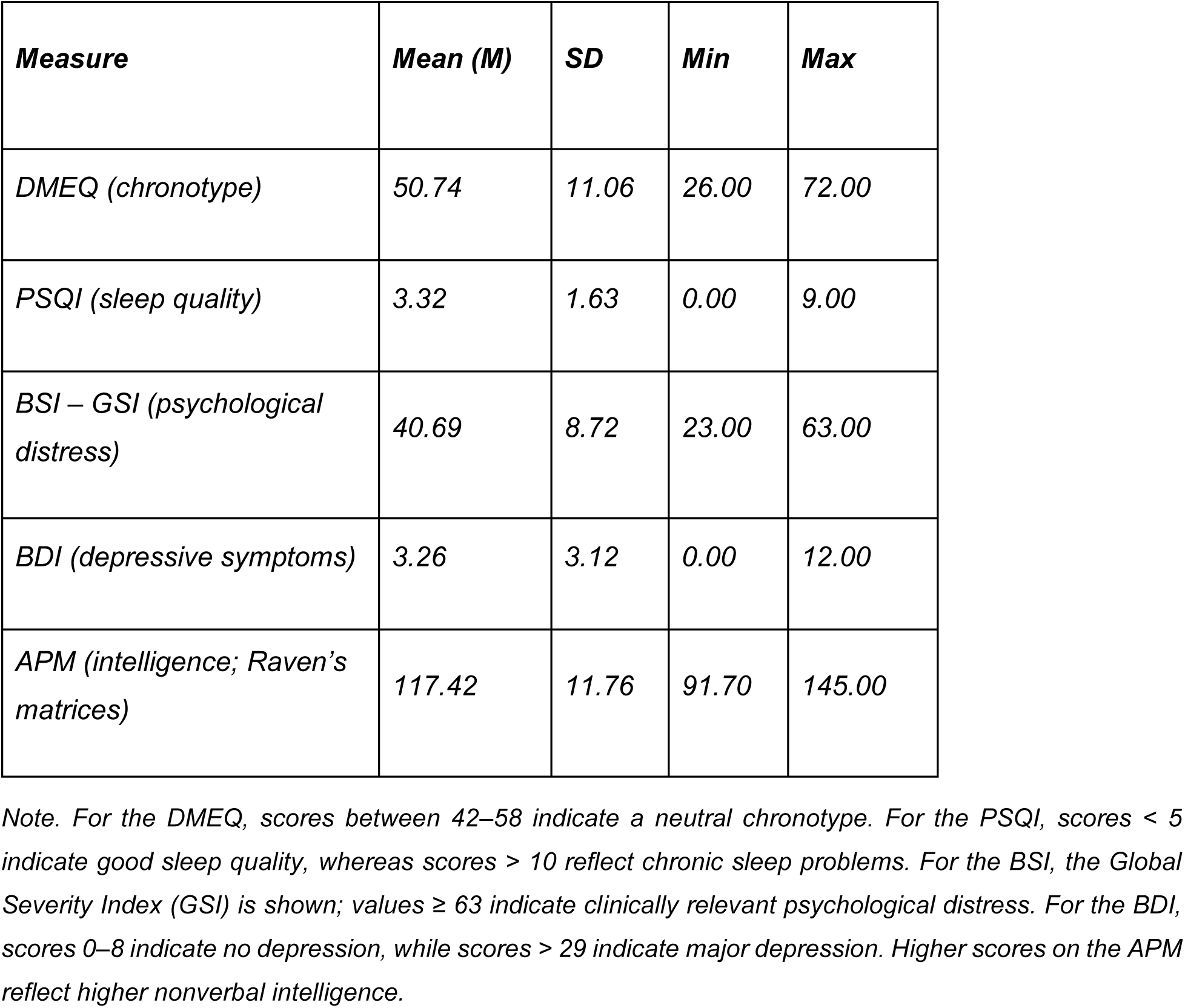
Descriptive statistics of questionnaire data.

**Supplementary Table 2.** Demographic sleep data.

**Demographic sleep data**
|  | **Adaptation night (M + SD)** | **Experimental Night (M + SD)** |
| --- | --- | --- |
| **TIB** | 478.60 ± 20.04 | 477.47 ± 15.34 |
| **TSP** | 455.73 ± 24.77 | 460.58 ± 30.50 |
| **WTSP** | 22.68 ± 34.14 | 11.98 ± 12.88 |
| **TST** | 433.05 ± 46.23 | 448.60 ± 32.50 |
| **WAFA** | 3.25 ± 8.11 | 5.60 ± 25.59 |
| **WASO** | 25.93 ± 37.37 | 17.58 ± 27.46 |
| **SOL** | 19.62 ± 11.64 | 11.28 ± 9.54 |
| **SE** | 90.49 ± 8.89 | 93.95 ± 5.94 |
| **PC_N1** | 10.64 ± 4.87 | 9.28 ± 4.40 |
| **PC_N2** | 41.21 ± 6.87 | 40.30 ± 7.29 |
| **PC_N3** | 30.97 ± 10.01 | 32.20 ± 10.03 |
| **PC_R** | 17.18 ± 5.05 | 18.22 ± 4.35 |

**Supplementary Table 3.**
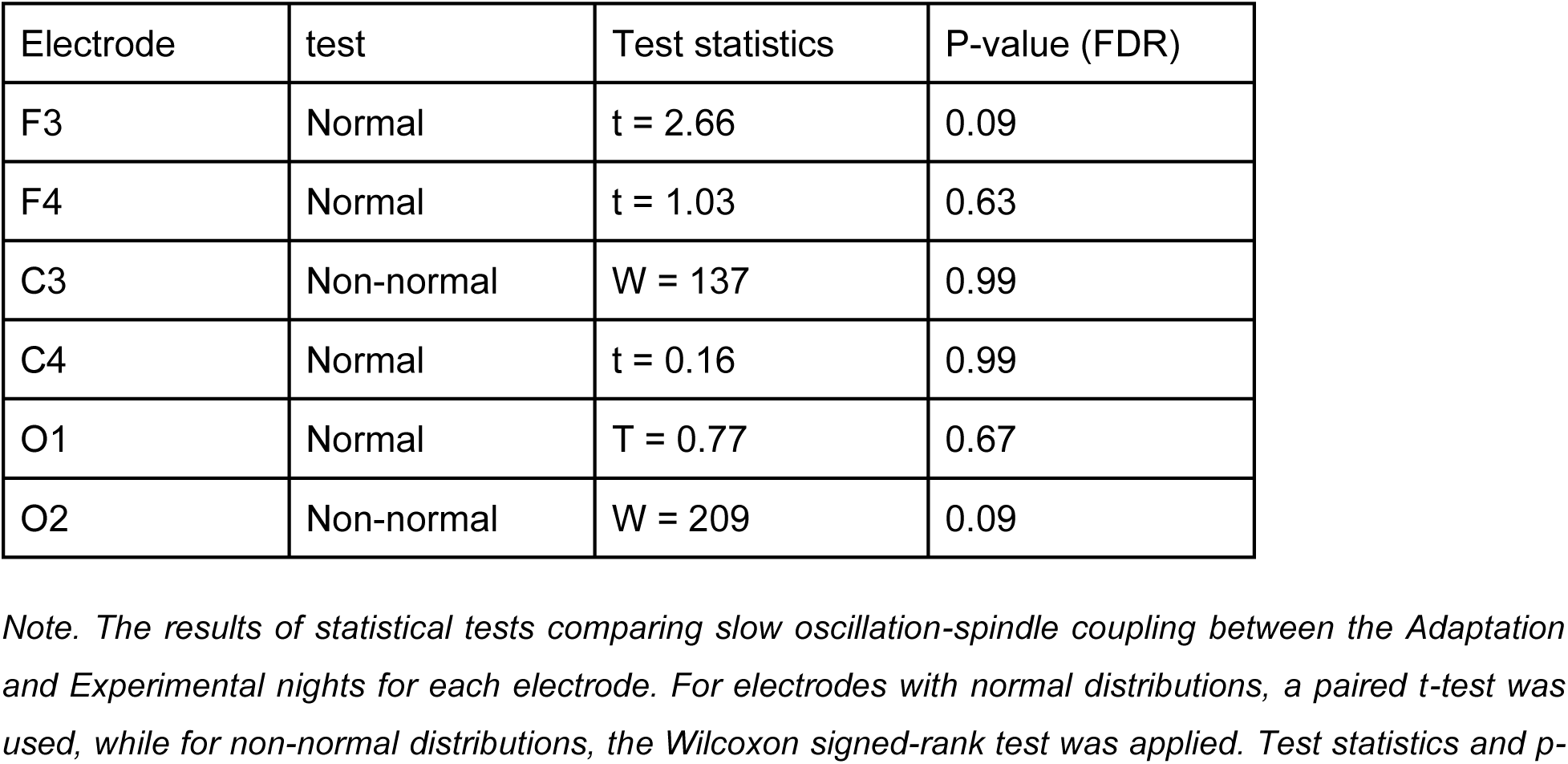
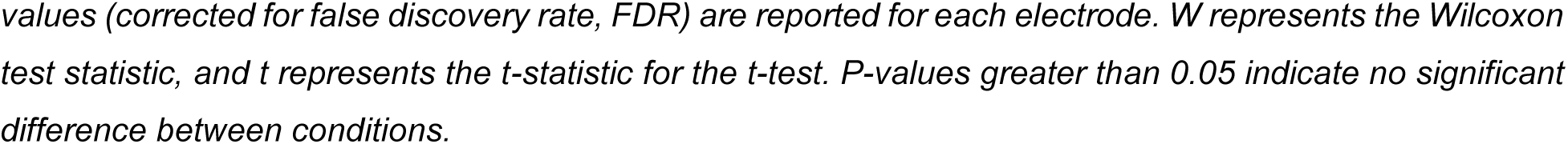
Statistical Tests for Slow oscillation-Spindle Coupling Across Electrodes.

| Electrode | test | Test statistics | P-value (FDR) |
| --- | --- | --- | --- |
| F3 | Normal | t = 2.66 | 0.09 |
| F4 | Normal | t = 1.03 | 0.63 |
| C3 | Non-normal | W = 137 | 0.99 |
| C4 | Normal | t = 0.16 | 0.99 |
| O1 | Normal | T = 0.77 | 0.67 |
| O2 | Non-normal | W = 209 | 0.09 |

*Note. The results of statistical tests comparing slow oscillation-spindle coupling between the Adaptation and Experimental nights for each electrode. For electrodes with normal distributions, a paired t-test was used, while for non-normal distributions, the Wilcoxon signed-rank test was applied. Test statistics and p-*
values (corrected for false discovery rate, FDR) are reported for each electrode. *W* represents the Wilcoxon test statistic, and *t* represents the *t*-statistic for the *t*-test. *P*-values greater than 0.05 indicate no significant difference between conditions.

**Supplementary Table 4.**
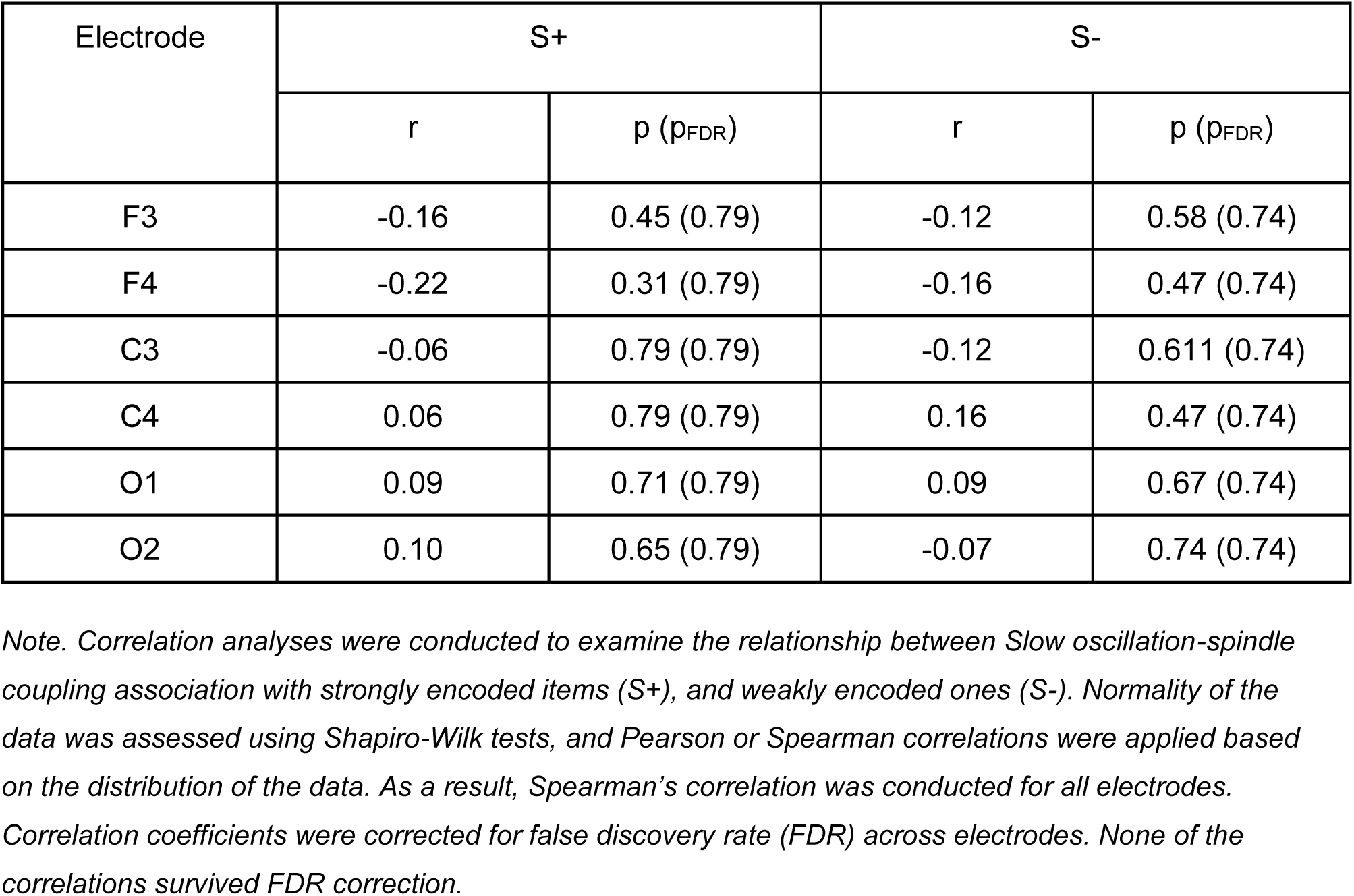
Correlation Analysis Results for Slow oscillation-Spindle Coupling and Behavioral Performance.

**Supplementary Figure 1.**
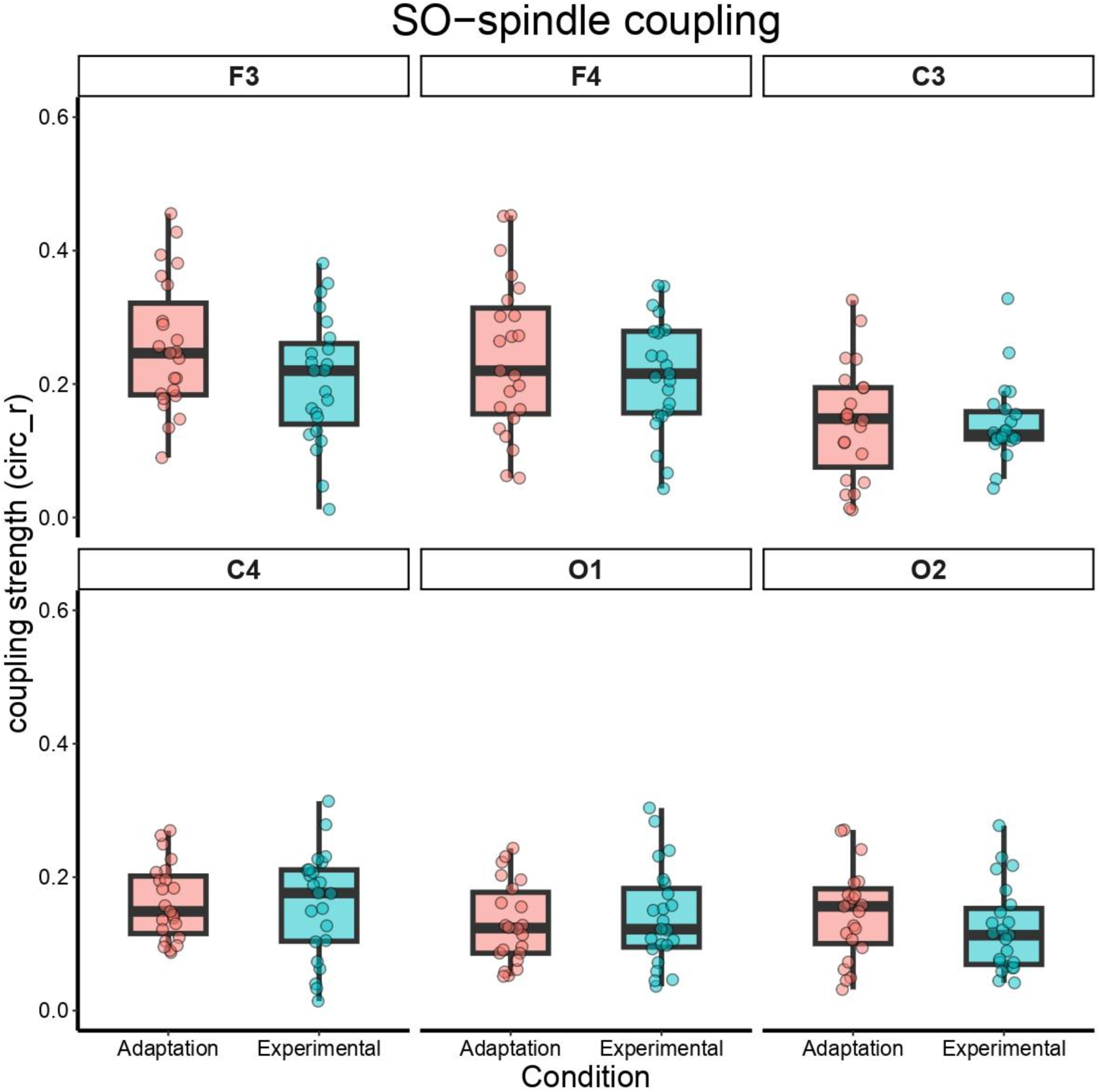
Difference in Slow oscillation-Spindle Coupling Between Adaptation and Experimental Nights Across All Electrodes. Coupling strength (measured as circular r) was compared between the Adaptation and Experimental nights across all electrodes. Boxplots were used to visualize the distribution of coupling strength for each condition, with individual subject data points overlaid to highlight within-subject variability. Statistical comparisons (paired t-test or Wilcoxon signed-rank test, depending on normality) revealed no significant differences between the conditions. The detailed statistical results can be found in Supplementary Table 3. Data points represent subjects’ means.

**Supplementary Figure 2.**
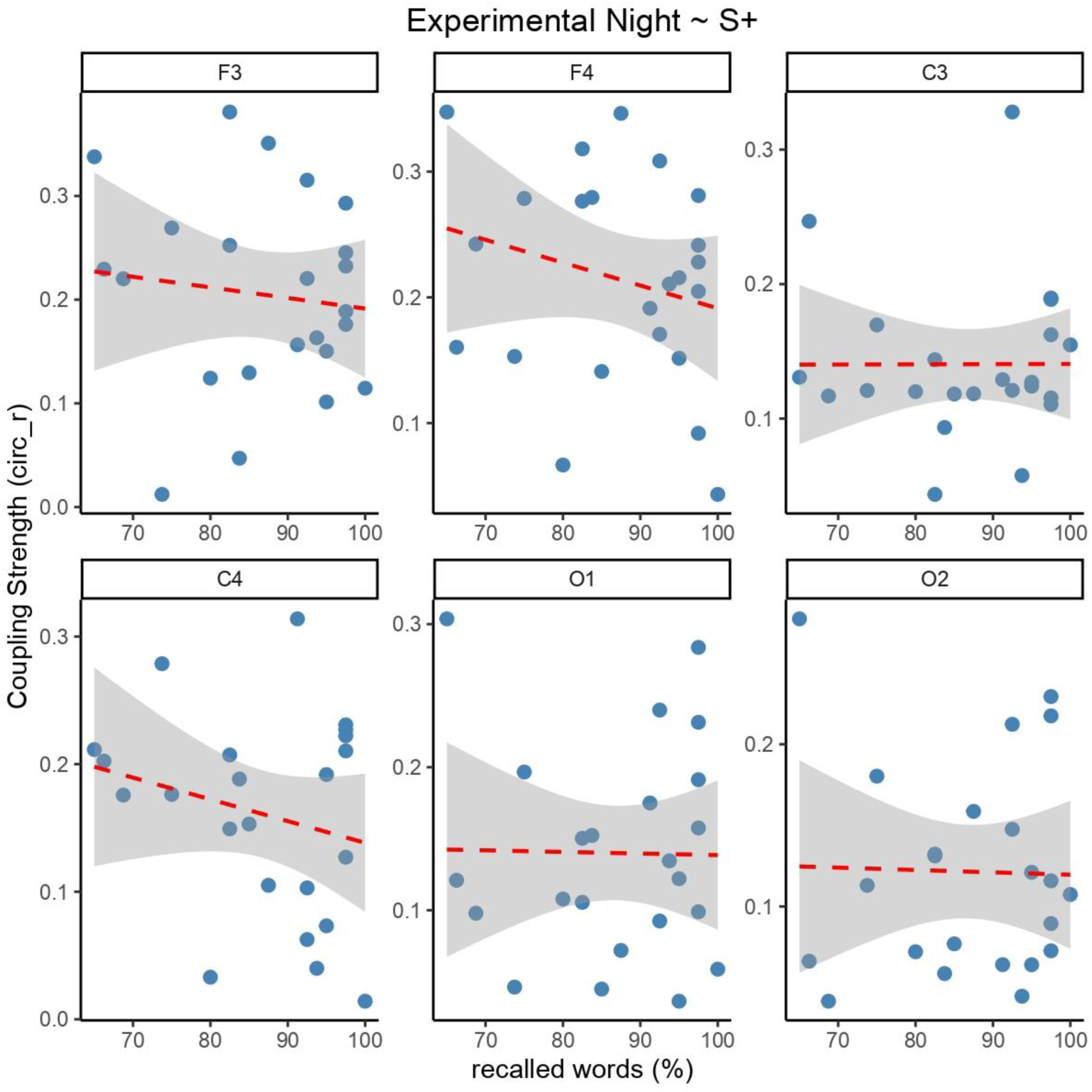
Correlation Between Slow oscillation-Spindle Coupling Strength and Recall of Strongly Encoded Items (S+) Across Electrodes. Spearman’s correlation analyses were conducted to examine the relationship between slow oscillation-spindle coupling strength and recall of strongly encoded items (S+) at each electrode. No significant correlations were found at any electrode. Each point represents data from an individual participant, and the red dashed line indicates the fitted linear regression line. The detailed statistical results can be found in Supplementary Table 4.

**Supplementary Figure 3.**
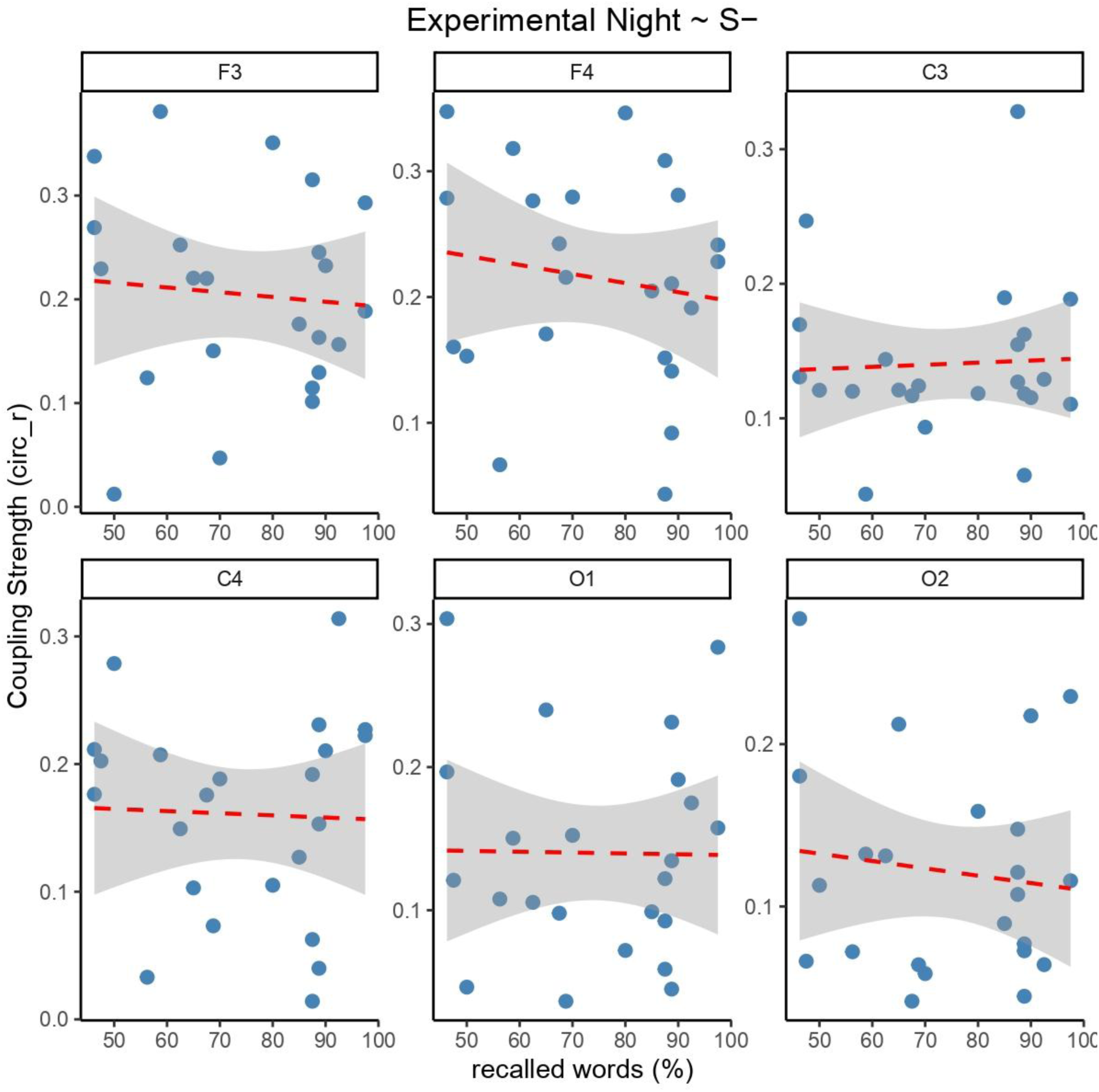
Correlation Between Slow oscillation-Spindle Coupling Strength and Recall of Weakly Encoded Items (S-) Across Electrodes. Spearman’s correlation analyses were conducted to examine the relationship between slow oscillation-spindle coupling strength and recall of weakly encoded items (S-) at each electrode. No significant correlations were found at any electrode. Each point represents data from an individual participant, and the red dashed line indicates the fitted linear regression line. The detailed statistical results can be found in Supplementary Table 4.

## Notes

### Competing Interest Statement

The authors have declared no competing interest.

### Summary of Updates

- Added a new section in the results to clarify the role of slow oscillation-spindle coupling. - added a new figuring comparing only SLEEP and WAKE groups. - Updated reaction times analysis. - Added demographic sleep data and psychometric data to supplements. - Revised wording in the whole manuscripts and added new sections in the discussion to discuss new results.

